# The human pupil and face encode sound affect and provide objective signatures of tinnitus and auditory hypersensitivity disorders

**DOI:** 10.1101/2023.12.22.571929

**Authors:** Samuel S. Smith, Kelly N. Jahn, Jenna A. Sugai, Ken E. Hancock, Daniel B. Polley

**Author notes:** Equal contribution (Samuel S. Smith, Kelly N. Jahn, Jenna A. Sugai). Department of Speech, Language, and Hearing, The University of Texas at Dallas, 1966 Inwood Road, Dallas, TX 75235.

## Abstract

Sound is jointly processed along acoustic and emotional dimensions. These dimensions can become distorted and entangled in persons with sensory disorders, producing a spectrum of loudness hypersensitivity, phantom percepts, and – in some cases – debilitating sound aversion. Here, we looked for objective signatures of disordered hearing (DH) in the human face. Pupil dilations and micro facial movement amplitudes scaled with sound valence in neurotypical listeners but not DH participants with chronic tinnitus (phantom ringing) and sound sensitivity. In DH participants, emotionally evocative sounds elicited abnormally large pupil dilations but blunted and invariant facial reactions that jointly provided an accurate prediction of individual tinnitus and hyperacusis questionnaire handicap scores. By contrast, EEG measures of central auditory gain identified steeper neural response growth functions but no association with symptom severity. These findings highlight dysregulated affective sound processing in persons with bothersome tinnitus and sound sensitivity disorders and introduce approaches for their objective measurement.

## Introduction

Damage or degeneration of peripheral sensory organs causes reduced sensitivity to environmental stimuli. It can also precipitate the opposite outcome: a hypersensitivity to environmental stimuli along with the perception of phantom stimuli that have no physical source in the environment. Sensory phantoms occur in all modalities but are more often heard than seen, tasted, felt, or smelled. Approximately 12% of adults hear an indefatigable phantom sound every day of their waking life. Tinnitus typically manifests as a continuous ringing, whooshing, roaring, or sizzling phantom sound that is often accompanied by a generalized sensitivity and discomfort with moderately intense environmental sounds^1,2^. Most evidence suggests that disinhibition in auditory processing centers of the brain is a proximal cause of tinnitus and sound sensitivity^3–7^. Central auditory disinhibition related to tinnitus and sound sensitivity can arise from normal aging^8,9^, as a compensatory response to hearing loss and auditory nerve degeneration^10–19^, from traumatic injury of the brain or cervical ganglia^20–22^, or simply from the abrupt cessation of prescription GABA agonists^23,24^. Whatever the source, the Excess Central Gain model posits that disinhibition has the knock-on effect of increasing the synchrony and activity rate of local excitatory neurons in silence (the basis of the phantom sound) as well as disproportionately strong responses to moderately intense sound (the basis of loudness hypersensitivity)^25–29^. As expected, boosting levels of central inhibition through direct activation protocols in animals or GABA-acting drugs in humans can mitigate tinnitus symptoms^30,31^.

In some cases, persons with tinnitus and sound sensitivity also present with an intense generalized aversion to sound, anxiety about sound exposure, social withdrawal, depression, and even suicidal ideations^32^. It is unclear how central gain in low-level auditory processing is related to the anxiety and mood disruptions that can accompany these disorders. One possibility is that the connection between elevated central gain and affective dysregulation may literally reflect abnormally strong coupling between auditory forebrain and limbic centers^33^. A recent animal study of auditory threat learning has shown that a selective plasticity in auditory corticoamygdalar projection neurons asymmetrically increased corticofugal spike-LFP coupling and sound-evoked activity in the lateral amygdala^34^, suggesting that increased neural gain in cortical projection neurons can be passed on to postsynaptic brain regions that regulate affective evaluation of valence and arousal. To this point, several neuroimaging studies in participants with tinnitus and sound sensitivity have identified abnormally strong coupling between auditory cortex and the amygdala, insula, anterior cingulate cortex, and medial prefrontal cortex^33,35–37^. Whereas psychoacoustic and low-level auditory assays were generally not correlated with extra-auditory features of sound aversion and psychological burden^38–43^, neuroimaging assays of enhanced coupling with extra-auditory networks have shown stronger correlations with individual differences in tinnitus and sound sensitivity severity^37,44^.

There are no objective measurements for the severity of tinnitus and sound sensitivity disorders. Instead, assessments rely on subjective self-report questionnaires, which can introduce vulnerabilities for malingering and false disability claims^39^ and offer less insight into the underlying causes and potential treatments. Here, we hypothesized that involuntary autonomic and behavioral responses may have untapped potential as non-invasive, objective markers of tinnitus and sound sensitivity severity that are relatively easily to implement in laboratory and clinical settings. Autonomic responses (e.g., pupil dilation and skin conductance) and involuntary facial movements provide a wealth of information into affective processing of valence and arousal in humans^45–48^ as well as other animals^34,49^. Although human studies have largely relied on visual stimuli, it is clear that speech, music and other sounds provide a rich medium for conveying valence and arousal cues^50–54^. This led us to ask whether emotionally evocative sounds elicit autonomic and facial responses that are modulated by perceived valence. We then evaluated whether these objective measures could identify a bias towards negative valence and hyper-arousal in persons with tinnitus and sound sensitivity that might more accurately predict the severity of sound aversion and psychological burden they report.

## Results

Of 196 adults recruited to our study, we identified 71 eligible participants with normal hearing thresholds who completed the full course of testing (**Figure S1A**). An experienced clinician assigned participants with chronic tinnitus and/or auditory hypersensitivity to the disordered hearing (DH, N=35) group and participants without tinnitus or auditory hypersensitivity to the neurotypical (NT, N=36) group. The distribution of age and hearing thresholds were closely matched between DH and NT groups. As expected, uncomfortable listening level thresholds in DH subjects were significantly lower than NT subjects (**Figure S1B-C**, description of statistical testing described in the figure legends throughout).

### Increased central gain is not correlated with tinnitus or sound sensitivity burden

The Enhanced Central Gain model posits that disinhibition in the auditory cortex and subcortical auditory structures produces hypersynchronized and hyperactive population activity among excitatory neurons in silence and hyperresponsivity to sounds of increasing physical intensity (**Figure 1A**). Although we could not directly measure the rate or synchrony of spiking, it was possible to measure auditory neural response gain through an approach that slowly increased and decreased the intensity of a 40 Hz amplitude modulated tone (**Figure 1B**, top). Scalp EEG recordings revealed a synchronized envelope following response (EFR) at 40 Hz that increased and decreased in amplitude with the change in sound intensity (**Figure 1B**, bottom). Gain (i.e., the output per unit step in input) could be explicitly measured as the slope of the neural growth function. As predicted by the Excess Central Gain model, neural activity grew significantly more steeply with increasing sound level in DH participants (**Figure 1C**).

**Figure 1.**
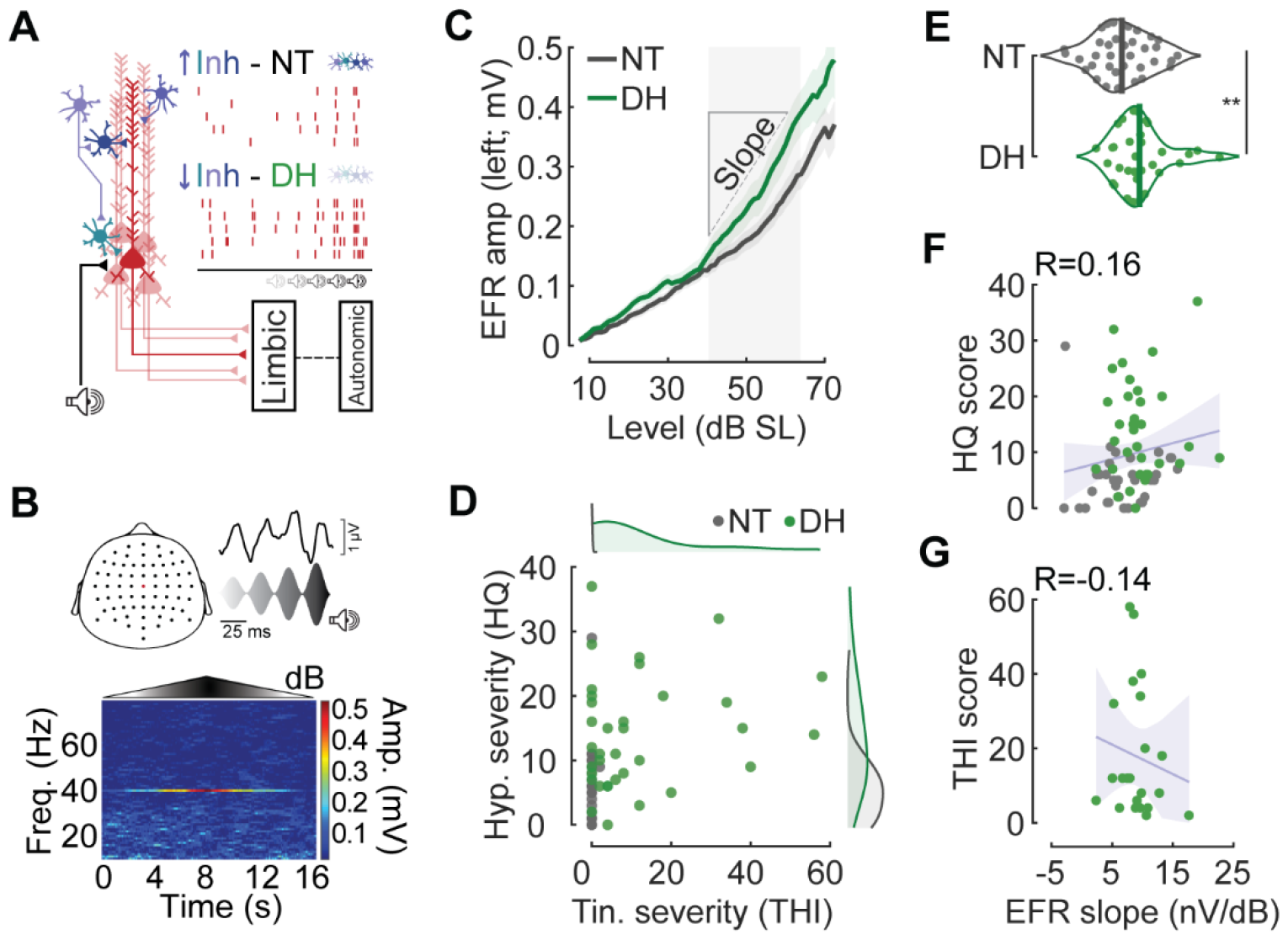
Increased central gain is not correlated with tinnitus or sound sensitivity burden. **A)** Cartoon denotes a stage of central auditory processing (e.g., the auditory cortex) with excitatory projection neurons (red) and inhibitory interneurons (cool colors). In this model, disinhibition of excitatory neurons promotes elevated, hypersynchronous firing in silence (the purported generator of the phantom sound) and a steeper growth in spiking with sounds of increasing intensity (i.e., excess central gain, the purported generator of loudness hyperacusis). Hyperactive auditory projection neurons feed into downstream centers of limbic processing and autonomic regulation but, as distal upstream precipitator, excess central gain is less predictive of individual differences in psychoaffective burden than autonomic affective markers. **B)** *Top:* Cartoon denotes the 64-channel array of scalp EEG electrodes and activity from a central electrode corresponding to the increasing intensity of a 40Hz amplitude modulated tone. Note that EEG amplitude is synchronized to the amplitude modulation rate. *Bottom*: Spectrogram plots the amplitude of synchronized EEG activity across frequencies and time as the amplitude modulated 2kHz tone slowly increases and decreases across a 70 dB range. Note the rise and fall of the 40Hz envelope following response (EFR) amplitude as a function of time/sound intensity. **C)** EFR growth as a function of sound intensity relative to the 2kHz audibility threshold measured for each participant (i.e., the sensation level, SL). NT and DH are neurotypical and disordered hearing participants, respectively. Central gain was measured as the change in neural response over a 25 dB change in sound level. **D)** Hyperacusis and tinnitus severity for all participants based on Hyperacusis questionnaire (HQ) and Tinnitus Handicap Index (THI) scores, respectively (N = 35/35 NT/DH). Circles denote individual participants. Marginal distributions for each group are shown as normalized density functions. All participants can provide a meaningful HQ score but only participants with tinnitus can provide a meaningful THI value. **E)** Central gain is significantly elevated in DH participants (two-sample t-test, p = 0.009, N = 36/33 NT/DH). Density functions display the central gain measure for each participant (individual circle) and sample mean (vertical lines). **F)** Central gain is not correlated with hyperacusis severity (Pearson R = 0.16, p = 0.19, N = 68). Shaded region denotes the 95% confidence interval. Solid line denotes linear fit. Circles denote individual participants. **G)** Central gain is not correlated with tinnitus severity (Pearson R = 0.13, p = 0.52, N = 22). Plotting conventions as per *e*. Note that THI values are limited to participants with tinnitus.

Tinnitus and sound sensitivity are heterogenous co-morbid conditions. The cohort of DH participants presented with varying degrees of sound aversion, distress, anxiety and depression related to their disorder, as identified by standard clinical questionnaires for hyperacusis and tinnitus (HQ and THI, respectively, **Figure 1D**). This variability provided us with a valuable opportunity to study the relationship between excess central gain and psychological burden. Although central gain was significantly increased at a group level in DH participants (**Figure 1E**, top), it was not correlated with individual differences in hyperacusis or tinnitus severity (**Figures 1F and 1G**, respectively).

As illustrated in the cartoon model (**Figure 1A**), an extension of the Excess Central Gain model posits that hyperactive, hypersynchronized, and hyperresponsive projection neurons in the central auditory pathway drive abnormally strong functional connectivity and auditory recruitment in downstream limbic networks. Limbic hyperresponsivity, in turn, would elicit abnormal autonomic response for sounds that would not otherwise be encoded as unpleasant or highly arousing. In essence, a failure to homeostatically regulate neural activity in auditory brain networks disrupts allostatic processes that balance autonomic and behavioral responses to stressors. In this model, excess central gain is a distal, upstream precipitator of limbic-autonomic dysfunction that more directly determines whether an individual feels utterly debilitated versus slightly annoyed by their tinnitus. As such, central gain would not be expected to be highly correlated with the tinnitus or hyperacusis burden reported in surveys, but a more direct marker of limbic or autonomic dysfunction would.

### Pupil-indexed affective processing is correlated with tinnitus and sound sensitivity burden

In constant light levels, pupil diameter provides an autonomic marker of brain-wide neuromodulator release^55,56^ that, depending on task design, can index executive load^57^, or affective valence and arousal^58^. To determine whether autonomic and behavioral evaluation of sound affect was biased towards negative valence and hyper-arousal in DH subjects, we measured pupil diameter and skin conductance while subjects listened to 6s audio clips from an established database of emotionally evocative sounds (**Figure 2A**)^59^. These sounds spanned a wide valence range, including sounds rated as highly unpleasant (screaming) to pleasant (music) (**Figure 2B**). On average, selected sounds smoothly tiled the valence range (**Figure S2A**) but were idiosyncratic at the level of individual participants, presumably reflecting their individual hedonic sound associations (**Figure S2B**). When individually ordered by valence rating, we found that DH subjects rated sounds as less pleasant overall, particularly relatively pleasant or neutral sounds, suggesting a complementary metric to the generalized sound aversion captured by subjective self-report questionnaires^60^ (**Figure 2C**).

**Figure 2.**
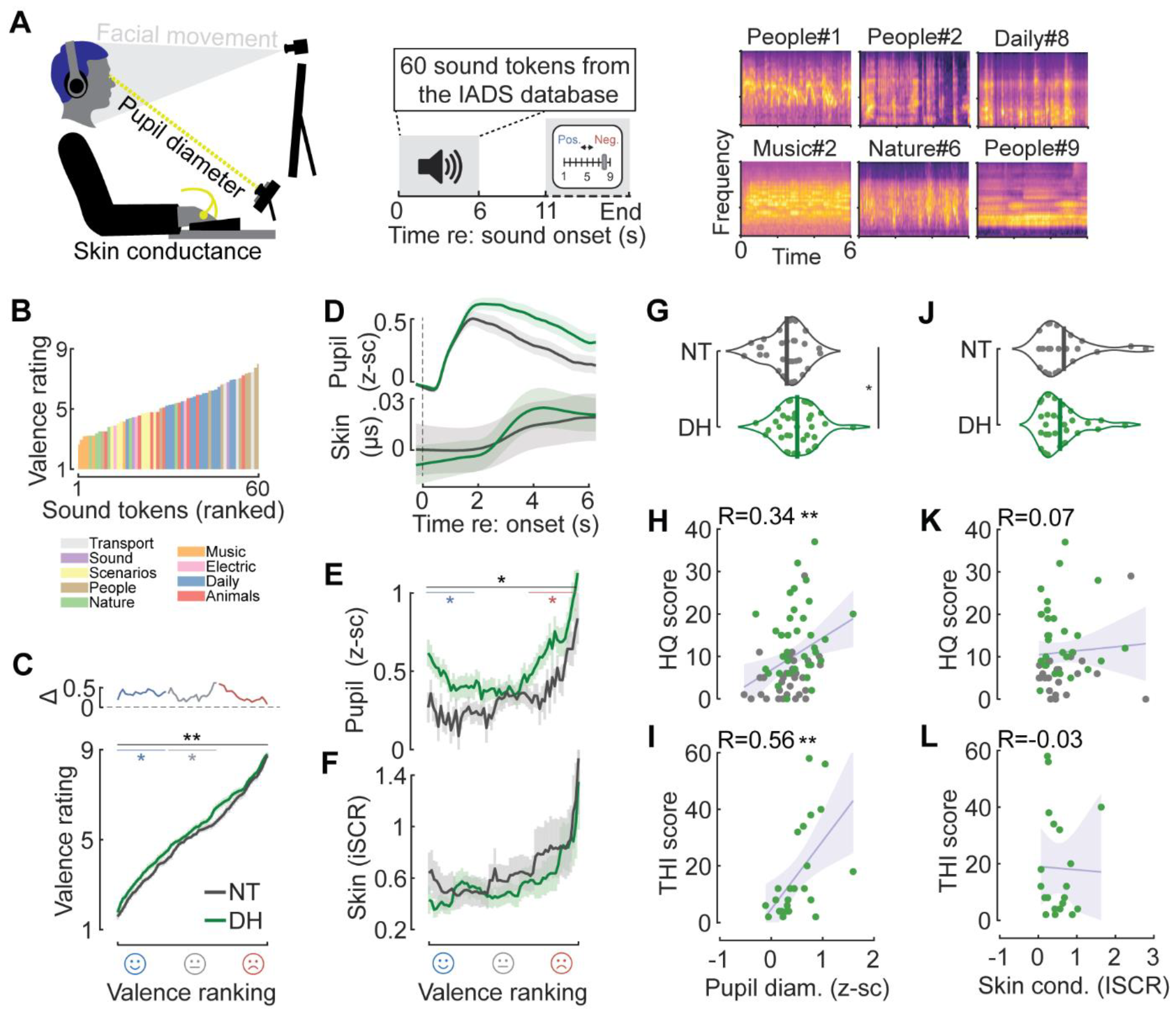
Pupil-indexed affective processing is correlated with tinnitus and sound sensitivity burden. **A)** Schematic of trial design and experimental setup for combined autonomic and behavioral evaluation of sound valence. Alongside are spectrograms of six representative sounds filtered with a gammatone filter bank. **B)** Mean behavioral valence rating of 60 sounds drawn from the IADS affective sound library (low/high scores indicate pleasant/unpleasant). Sounds are shown rank ordered and color-coded by labeled category. **C)** Mean individually rank-ordered valence ratings demonstrate a significant bias overall towards unpleasant ratings in DH participants (Wilcoxon rank sum, p = 0.005, N = 36/35 NT/DH). When discretized into pleasant, neutral, and unpleasant categories (blue/gray/red, respectively), valence ratings were significantly elevated for pleasant and neutral sounds (asterisks, Wilcoxon rank sum, p = 0.026 and p = 0.048, respectively). **D)** Mean ± SEM pupil diameter and skin conductance for all sounds during the 6s stimulus period. Responses are grouped by NT and DH participants. **E)** Sound-evoked pupil dilations were larger overall in DH participants (two-sample t-test, p = 0.03, N=32/35 NT/DH). Pupil dilations were significantly increased for pleasant and unpleasant sounds (two-sample t-tests, p = 0.01 and 0.03, respectively) in DH compared to NT subjects. **F)** Skin conductance did not differ overall (two-sample t-test, p = 0.59, N = 24/30 NT/DH) or as a function of valence (two-sample t-tests, p > 0.5 for each valence category) in DH compared to NT subjects. **G)** Density functions display the sound-evoked pupil response averaged for each participant (individual circle) and sample mean (vertical lines). **H)** Pupil size is correlated with HQ score (Pearson R = 0.33, p = 0.006, N = 66). Shaded region denotes the 95% confidence interval. Solid line denotes linear fit. Circles denote individual participants. **I)** Pupil size is correlated with THI score (Pearson R = 0.56, p = 0.005, N = 24). **J)** Density functions display the skin conductance response averaged for each participant (individual circle) and sample mean (vertical lines). **K)** Skin conductance is not significantly correlated with HQ score (Pearson R = 0.07, p = 0.63, N = 53). Shaded region denotes the 95% confidence interval. Solid line denotes linear fit. Circles denote individual participants. **L)** Skin conductance is not significantly correlated with THI score (Pearson R = -0.03, p = 0.92, N = 20).

We observed that emotionally evocative sounds elicited pupil dilations within approximately 0.5s followed by a slower increase in sound-evoked skin conductance (**Figure 2D; Figure S3**). Pupil dilation scaled with self-reported sound valence across all participants but was significantly greater overall in DH subjects, indicating a hypersensitized autonomic response to both pleasant and unpleasant sounds (**Figure 2E**). Sound-evoked changes in skin conductance also scaled with self-reported valence but did not differ between groups (**Figure 2F**). Sound-evoked pupil dilation was significantly elevated in DH participants (**Figure 2G**) but, unlike central gain, was significantly correlated with individual differences in self-reported hyperacusis burden (**Figure 2H**) and tinnitus burden (**Figure 2I**). Average sound-evoked skin conductance did not differ between groups and was not correlated with tinnitus or hyperacusis severity (**Figure 2J-L**).

### Pupil hyper-responsivity is specific to affective processing

These findings support our hypothesis that affective sound encoding is disrupted in persons with debilitating tinnitus and sound sensitivity, which can be measured through autonomic markers of affective arousal such as pupil dilation. Two negative control experiments support the assertion that pupil-indexed hyper-arousal was specific to affective processing. First, pupil dynamics in DH and NT participants were indistinguishable when entrained to periodic changes in light intensity (**Figure S4A-B**). Second, we measured pupil-indexed listening effort in a multi-talker speech intelligibility task (**Figure S4C**) and observed reduced behavioral accuracy and increased pupil diameter at a more challenging signal to noise ratio (**Figure S4D**). Importantly, behavioral accuracy and pupil-indexed listening effort were equivalent between NT and DH participants, confirming that DH subjects do not have widespread deficits in challenging listening tasks^61^ or global autonomic regulation, but rather a more specific behavioral and autonomic aversion to the affective quality of sound (**Figure S4E-F**).

### Subtle facial movements identify auditory valence and hearing disorder severity

To identify additional windows into auditory affective processing that might complement or extend upon pupil dynamics we turned our attention to subtle and rapid movements of the face. Facial movements have a long history in human emotion research, though these studies have almost exclusively relied on visual stimuli to convey affective cues^62^. To establish whether emotionally evocative sounds would also produce facial movements related to affective valence and arousal, we quantified high-resolution facial videography recordings that were performed in tandem with the pupil and electrodermal recordings (**Figure 3A**). We developed an analysis pipeline to localize changes in facial texture and observed that sounds do elicit subtle and rapid facial movements (**Figure 3B**). For example, in one instance an unpleasant “buzz” sound elicited a tightening of the muscles around the eyes and forehead (**Figure 3C**, left), whereas a more pleasant sound elicited a longer latency movement of the mouth and jaw (**Figure 3C**, right). Amongst this spatiotemporal complexity, we found stereotyped patterns that were associated with individual valence ratings. Pleasant sounds evoked increased movements in the area of the zygomaticus major, a muscle involved in shaping the corners of the lips (**Figure 3D**, left). Unpleasant sounds triggered increased movements globally but were particularly robust around the eyes and brows (**Figure 3D**, middle).

**Figure 3.**
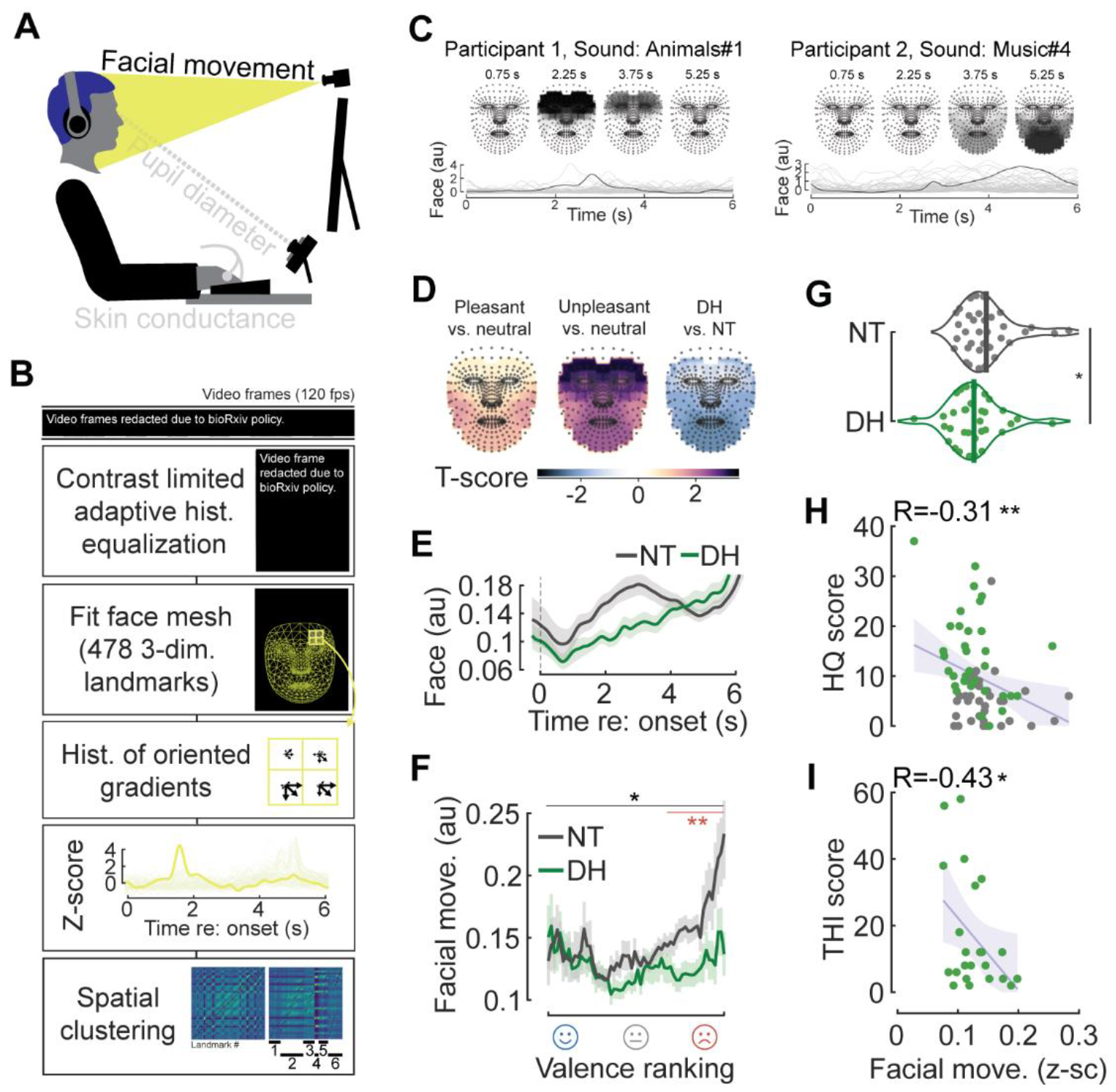
Subtle facial movements identify auditory valence and hearing disorder severity. **A)** Facial videography experimental setup. **B)** Videos were processed with a custom analysis pipeline incorporating a face-mesh mapping and subsequent pixel quantification (see Methods). **C)** Single trial facial movement amplitudes in two participants illustrate differences in amplitude and time course for an unpleasant (*left*) and pleasant (*right*) sound. **D)** T-score contrast maps of differences in facial movements for unpleasant vs neutral (*left*), pleasant vs. neutral (*middle*), and for NT vs. DT participants (*right*) during the 6s stimulus period. **E)** Mean ± SEM facial movement for all sounds during the 6s stimulus period. Responses are grouped into NT/DH. **F)** Facial reactions were reduced overall in DH participants (two-sample t-test, p = 0.046, N = 36/33 NT/DH) and were significantly reduced for unpleasant sounds (two-sample t-test, p = 0.003) in DH compared to NT subjects. **G)** Density functions display sound-evoked facial movements averaged for each participant (individual circle) and sample mean (vertical lines). **H)** Facial movement is negatively correlated with HQ score (Pearson R = -0.31, p = 0.0096, N = 68). Shaded region denotes the 95% confidence interval. Solid line denotes linear fit. Circles denote individual participants. **I)** Facial movement is negatively correlated with THI score (Pearson R = -0.42, p = 0.042, N = 23).

Like the pupil, we hypothesized that sound-evoked facial expressions would convey downstream limbic dysfunction in individuals with tinnitus and sound sensitivity. This prediction was upheld but, interestingly, in the opposite direction of changes in pupil dilation. Whereas facial movement amplitude scaled with valence in NT participants, DH participants exhibited a generally blunted affect that did not change across self-reported valence (**Figure 3E-G**). Importantly, reduced facial movements were significantly correlated with individual differences in sound sensitivity burden (**Figure 3H**) and tinnitus burden (**Figure 3I**). Together, these results show that the face encodes not only sound affect but can also provide a window into the intense aversion and lifestyle burden that can accompany tinnitus and sound sensitivity.

### Combined measures of sound affect best determine psychological burden

Here, we have identified neural (**Figure 1C**), autonomic (**Figure 2E**), voluntary behavioral (**Figure 2C**), and involuntary behavioral (**Figure 3F**) measures that distinguish DH and NT subjects at a group level. A major goal for research on sensory disorders is to identify and refine a set of objective measurements – akin to what tumor imaging and biopsy data provide for oncologists or EEG measures of epileptiform activity provide to neurologists – that inform future care providers about the subtype and severity of the sensory disorder, the likelihood to benefit from a particular treatment, or whether they have demonstrated an improvement subsequent to treatment. While some measurements have been developed to identify whether an individual has tinnitus, they are insensitive to severity and lifestyle burden^39,63,64^. Here, we found that pupil dilation and facial reactions both demonstrated a correlation with symptom severity. As a next step, we determined whether these measures were redundant and how accurately they could predict an individual’s self-reported symptom severity.

We incorporated all measures where DH and NT cohorts were significantly different at a group level (pupil diameter, facial movement, behavioral valence rating, central gain) as well as potential co-variates of lifestyle burden (age) in an elastic net regression analysis, developed to be robust to superfluous predictor variables (see Methods). For the severity of self-reported sound sensitivity, a cross-validated fitting procedure settled on an optimal model that combined measures of affective responses – pupil dilation, facial movement, and behavioral valence rating (**Figure 4B**). This model was significantly correlated with hyperacusis questionnaire scores and was moderately accurate in predicting the hyperacusis index score compiled across NH and DT participants (R = 0.6/R^2^ = 0.36, **Figure 4C**). When the same approach was applied to tinnitus, age, we noted that central gain, and behavioral valence rating were minimally weighted, a moderate weight was afforded to pupil dilation, but facial reactivity was heavily weighted (**Figure 4D-E**). This model was highly correlated with self-reported severity and exhibited fairly high accuracy in predicting individual tinnitus burden scores (R = 0.78/ R^2^ = 0.61, **Figure 4D-F**). These findings demonstrate that pupil and facial reactivity to emotionally evocative sounds capture distinct features of generalized aversion, distress and anxiety that can accompany tinnitus and sound sensitivity. Although autonomic measures and affective processing have rarely been considered as biomarkers for these conditions, predictions of hyperacusis and tinnitus severity scores were significantly poorer when they were left out and the optimal model instead relied on neural and voluntary behavioral measures (likelihood ratio test, p<0.001).

**Figure 4.**
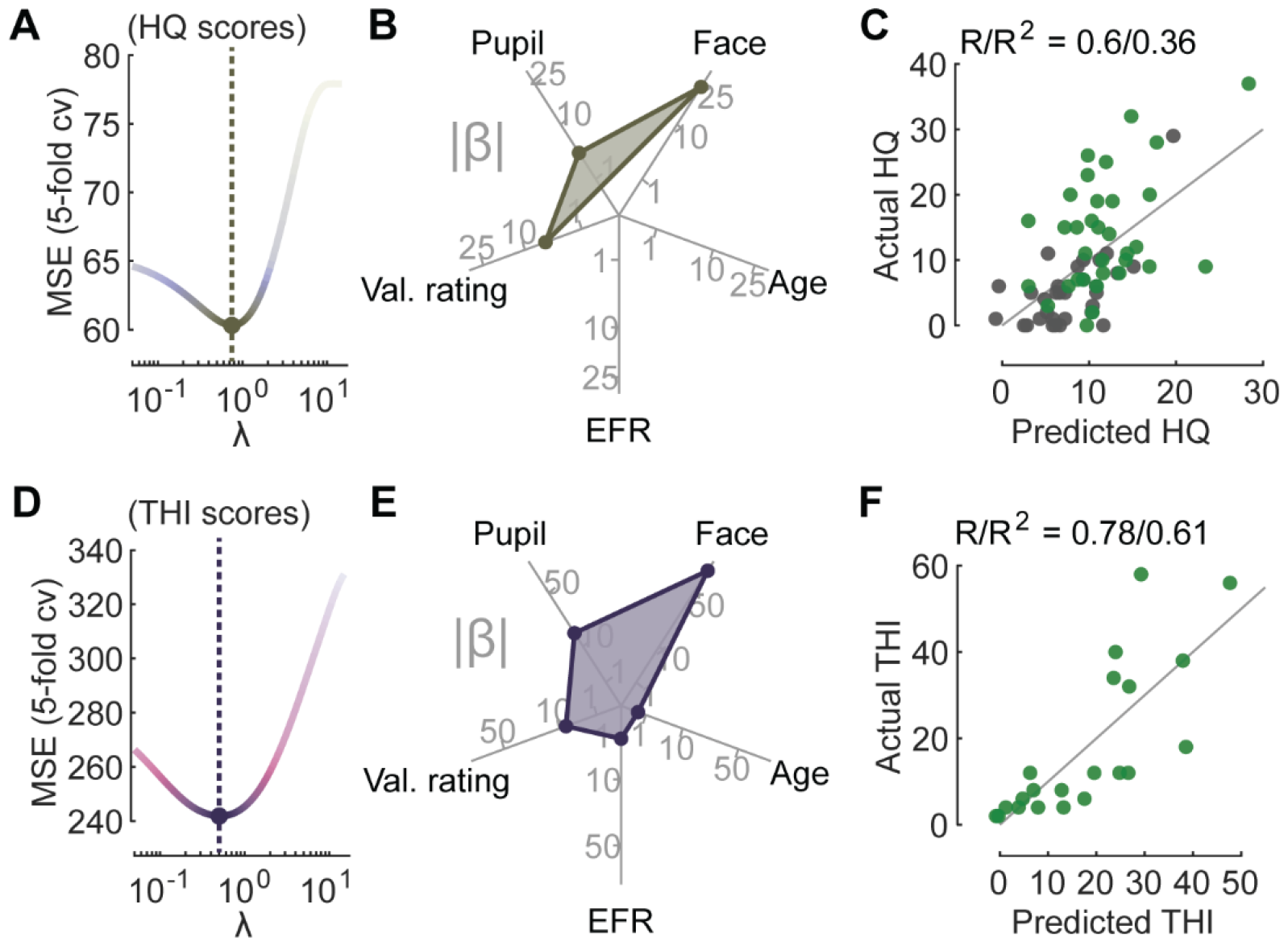
Combined measures of sound affect best determine psychological burden. **A)** An elastic net regression model was fit to individual hyperacusis severity scores (HQ scores). A tuning parameter, λ, controls the extent to which the coefficients contributing least to predictive accuracy are suppressed. **B)** A radar plot displays the coefficient weight (i.e., |β|) in the optimal model of HQ score (filled shape), The optimal model with minimal cross-validated error retained non-zero coefficients for pupil, facial movement, and valence ratings. **C)** The HQ scores predicted via elastic-net regression versus participants’ actual scores. **D)** Plotting conventions as per *A*, for tinnitus burden (THI) scores. **E)** Plotting conventions as per *B*, for THI scores. Predictors from the optimal elastic-net featured physiological measures of sound affect (face, pupil) as the highest weighted coefficients. **F)** Plotting conventions as per *C*, for THI scores.

## Discussion

### Extending upon the Excess Central Gain model of tinnitus and sound sensitivity

Pioneering neuroimaging studies in human subjects with tinnitus and reduced sound level tolerance identified auditory hyper-responsivity in the inferior colliculus and auditory cortex as the proverbial “ghost in the machine”^65^, i.e., a biological substrate for these perceptual disorders^42,66^. Subsequent findings in human subjects and animal models highlighted the juxtaposition of normal or even hyper-responsive event-related potentials arising from later stages with reduced sound-evoked potentials arising from the auditory nerve^12,15,67^ Findings of this ilk laid the foundation for the Enhanced Central Gain theory for tinnitus and sound sensitivity, wherein central auditory circuits leverage disinhibition to compensate for reduced bottom-up input. In so doing, vulnerabilities for hyper-synchrony, hyperactivity, and abnormally steep sound intensity growth functions are introduced that generate phantom sounds and reduced tolerance of moderately intense sounds^25,26^. Here, we used 40Hz amplitude modulation to enhance the relative contribution of the auditory cortex to the EFR^68^ and derived the instantaneous amplitude as the sound level swept up and down across a 70 dB range to provide a direct demonstration of enhanced central gain (i.e., increased neural output per unit step in sound input) in participants with tinnitus and reduced sound level tolerance.

Disinhibition and enhanced central gain at later stages of the auditory pathway may generate the sensory qualities of the phantom sound and disproportionate loudness but – in and of itself – is unlikely to account for the attentional and affective sequelae of tinnitus and sound sensitivity. Central gain does not account for why the phantom sound effortlessly recedes from conscious awareness when not attended to for some but is irrepressible for others. Likewise, central gain does not account for why internally and externally generated sounds a mild nuisance for some but are irritating, overwhelming, and anxiety-producing in others? That central gain does not account for these qualities does not mean that phenomenological models featuring auditory nerve degeneration and enhanced central gain are wrong, but rather that they are incomplete^69^.

Excitatory neurons from primary and higher-order auditory cortex project widely throughout prefrontal cortex executive control networks and limbic brain centers in neocortex and amygdala^70–72^. While relatively complex multi-regional feedback loops^73^, altered perceptual inference mechanisms^74^, or stochastic resonance^75^ may account for the attentional and affective components of tinnitus and sound sensitivity disorder, a more parsimonious explanation that directly builds upon the Excess Central Gain model is simply that hyperactive auditory efferent projections into extra-auditory executive and limbic networks produce functional hyperconnectivity and knock-on dysregulation in these networks, and this dysregulation more directly accounts for the attentional and affective components of these disorders.

Support for this model can be found in reports of enhanced auditory-limbic network connectivity in animals and humans^33,35–37^, though more detailed accounts of dysregulated spontaneous and sound-evoked processing in these regions are needed. Importantly, indirect but objective indices of affective and executive disruptions do not require specialized, costly, and low-throughput brain imaging systems. Pupil diameter, eye position, and involuntary facial reactivity have a well-established relationship with executive and affective processing and task designs to probe these relationships in neurotypical and neurodivergent populations have been the subject of study for many decades^49,58,76^. Here, we adapted pupil and facial assays of affective processing to the auditory modality and demonstrated their improved ability to account for the affective component of tinnitus and sound sensitivity disorder. While pupil and behavioral measurements reported here and elsewhere did not suggest deficits related to executive load or listening effort^61^, the attentional component of tinnitus disorder awaits more direct interrogation with paradigms that probe the persistence of attention to invariant sensory features with varying implicit significance.

### Limitations of the current study

One general limitation of observational research on human sensory disorders relates to uncertainty about the underlying precipitators of tinnitus and sound sensitivity in our subjects. Hearing loss, age, and certain medications can directly impact central gain and tinnitus^8-24^, motivating us to strictly screen and match these variables in NT and DH cohorts to control for a confounding influence. Nevertheless, the underlying cause and duration of tinnitus and sound sensitivity in our DH cohort is unknown and may contribute to the heterogeneity reported here. Another limitation is that the EEG, pupil, and facial measures described here provide little insight into their underlying generators in the central and autonomic nervous systems. We leveraged the excellent temporal resolution of EEG to derive measures of central gain that would be impossible with measures like fMRI but acknowledge that non-invasive imaging approaches could significantly extend and enrich the observations reported here. Another consideration is that while optimized multi-variate models featuring these objective markers of affective processing captured a sizeable fraction of the variance in clinical outcome measures and clearly outstripped other predictors studied here, these linear models fall short of providing the sensitivity and selectivity required of a diagnostic measure. One important point to consider here is that predictive accuracy cannot exceed the internal noise (e.g., test re-test reliability) and internal validity of the outcome measure. Whether because tinnitus and sound sensitivity are inherently dynamic disorders that ebb and flow over time or because the structure of the questionnaires elicit variable answers, the upper bound of the predictive accuracy reported here is capped by the reliability of the outcome measures themselves^77^. One of the underlying motivations for our study was to develop alternatives to self-report questionnaires, not only because of their implicit subjectivity but also because their validity as an instrument to probe the psychological burden arising from their tinnitus and hyperacusis independent of their broader mental state is not certain^78,79^. We propose that the predictive accuracy will be further improved by distilling the affective sound features used in the test battery, and combining these refined stimulus sets with ecological momentary assessment approaches that embrace the inherent dynamics of these disorders instead of reducing these subjects to a single measure of central tendency.

### Broader implications for clinical research on sensory disorders

The affective biomarkers described here were non-invasive and could be measured without specialized equipment, suggesting a promising approach for complementing subjective self-report questionnaires in clinical settings. These measures could prove to be useful biomarkers for other neurological disorders where auditory aversion is a prominent clinical feature, particularly for neurodevelopmental disorders such as autism spectrum disorder, where participant age or language impairment may preclude objective assessments based on questionnaires^80,81^. Objective physiological or neuroimaging biomarkers have been essential for the development of effective treatments for epilepsy, stroke, and other neurological conditions. Better established objective biomarkers for symptom severity in heritable and acquired sensory disorders will accelerate the pace of identifying therapies for these conditions^82^.

## Supplemental Information

A supplementary document consisting of four supplemental figures is available to download.

## Acknowledgements

This research was supported by research awards from the National Institute on Deafness and Other Communication Disorders to D.P. (P50DC015857) and K.J. (K01DC019647). Additional support for K.J. was provided by an ASHFoundation New Investigators Research Grant.

## Author contributions

K.J. and D.P. conceived the project and designed the experiments. J.S. collected the data using software programmed by K.H and supported by K.J. and S.S. Data analysis was led by S.S. with contributions from K.J. Figure preparation and manuscript writing was performed by S.S. and D.P. with input from all authors.

## Declaration of Interests

The authors declare no competing interests.

## Methods

### Experimental model and subject details

All procedures were approved by the Mass General Brigham Institutional Review Board.

#### Participants

Data are from 71 adults between 19 and 60 years of age (mean age = 33.9 years, 42 females) that were fluent in English. Participants were recruited as part of a larger study through flyers, word of mouth, and by posting to the Mass General Brigham participant recruitment website. As detailed in **Figure S1A**, screening and grouping of the 196 potentially eligible participants were performed by licensed clinicians. Participants were required to have normal hearing (unremarkable otoscopic evaluation and air conduction thresholds for tones 0.25 to 8 kHz ≤ 25 dB HL, 75 participants excluded). Participants were required to have normal cognitive function (telephone Montreal Cognitive Assessment, t-MOCA ≥18). We assessed mental health status and an ability to tolerate sound stimuli used in our experiments based on their response to question 24 in Beck’s Depression Inventory, question 23 of the Tinnitus Reactivity Questionnaire, and question 23 of the Sound Reactivity Questionnaire (10 participants excluded). Of the remaining participants, 16 were lost to follow-up after the initial orientation and four were excluded for not completing the full course of testing.

Participants were assigned to neurotypical or disordered hearing groups by experienced clinicians based on their responses to an open-ended questionnaire and clinical evaluation. Participants were excluded if they had catastrophic tinnitus (> 77 on Tinnitus Handicap Index, THI, 1 participant excluded), intermittent tinnitus (14 participants excluded), or inconsistently reported having tinnitus across questionnaires (4 participants excluded). Of the remaining participants, those who did not report sound sensitivity, nor intermittent or chronic tinnitus were classified as neurotypical (N=36, 25 females). The remaining subjects were assigned to the disordered hearing (DH) group (N=35, 17 females) based on their clinical evaluation of chronic tinnitus and/or abnormal sound sensitivity. Among these participants, we found that EEG data (N=2) and skin conductance recordings (N=17) were not usable on account of poor electrode contact. We excluded pupil data (N = 4) and facial movement (N = 2) due to inordinately high rates of blinking or failed tracking.

### Method details

Participants performed psychophysical and questionnaire assessments remotely, between two in-lab sessions (approximately 3 hours each).

#### Patient reported outcome measures

The psychoaffective burden of tinnitus and hyperacusis symptoms was assessed with the Hyperacusis Questionnaire (HQ)^83^ and the Tinnitus Handicap Inventory (THI)^84^. Both NT and DH participants can provide meaningful questionnaire responses to the HQ, which focuses on discomfort and aversion experienced with environmental sounds. By contrast, only participants who experience tinnitus can provide meaningful responses to the THI, which focuses on the lifestyle burden associated with phantom auditory percepts.

#### Pure tone audiometry and uncomfortable loudness level testing

Prior to EEG recordings, air conduction thresholds were measured for each ear with insert earphones (EarTone-3A) for pure tones ranging from 0.25 – 8 kHz in octave intervals using the modified Hughson-Westlake procedure. Uncomfortable loudness level (ULL) assessment was also performed for each ear just prior to the EEG session as well as during the remote tablet-based testing. Prior to the EEG session, ULL was determined with the Contour Test of Loudness Perception^85^, which presented three amplitude modulated 2kHz tones (200ms duration, 5ms raised cosine onset and offset ramps) beginning 5-dB below their 2 kHz hearing threshold and ascended in 5-dB steps until the participant rated the sound as “Uncomfortably Loud”. Participants completed four runs and the median intensity level across the four runs determined the demarcation of all seven loudness categories. Tablet-based testing was performed with a calibrated tablet computer and circumaural headphones (Bose AE2). For tablet-based testing, participants dragged a virtual slider to adjust pure tone sound intensity to the point where sound was judged to be uncomfortably loud, as described previously^77^. The ULL was the average across three repetitions.

#### Electroencephalography (EEG) measurements of central gain

Participants were seated in a reclining chair within a sound-treated booth and watched a movie of their choice with the volume muted and subtitles on. Continuous audio and EEG monitoring ensured that participants remained in an awake, restful state for the duration of testing. EEG recordings were performed with a 64-channel array of scalp electrodes and insert earphones (EarTone 3A) connected to an electrically isolated digitizer and signal processor (BioSemi ActiveTwo, Cortech Solutions Inc.). Envelope following responses (EFRs) were measured in response to auditory stimuli delivered to the left ear via insert headphones (EarTone-3A). Stimuli were sinusoidal amplitude-modulated tones with a carrier frequency of 2 kHz, a modulation rate of 40-Hz, and 100% modulation depth. Stimulus delivery (sampling rate: 100 kHz) and data acquisition (sampling rate: 8192 Hz) were coordinated and aligned through custom LabVIEW applications. The stimulus was repeated 160 times with alternating polarity of the carrier signal. Stimulus intensity was continuously ramped up (8 seconds) and down (8 seconds) over a 70 dB range, corresponding to a rate of intensity change of 8.75 dB/second. The upper limit of the intensity range was initially set to the median sound rated 6, “Loud, but comfortable” with additional downward adjustments made upon subject request.

#### Affective sound evaluations

Participants listened to 60 emotionally evocative environmental sounds from the original International Affective Digitized Sounds (IADS) corpus and the extended IADS-E corpus^59,86^. The IADS corpus contains 167 naturally occurring sounds that have been extensively validated for quantifying differences in affective reactions. For the present study, a subset of sixty IADS stimuli were chosen to span previously established valence categories^58,86,87^. Stimuli were presented binaurally through calibrated circumaural headphones (Bose AE2) in random order at a root-mean-square level of 75 dB SPL. A single trial consisted of a 5-second pre-stimulus baseline period, a 6-second stimulus, a 5-second silent post-stimulus period, and 30 seconds for the participant to self-report their behavioral response to the stimulus and for the physiological responses to return to baseline. Behavioral evaluations asked participants to rate the valence, arousal, and loudness of the preceding sound with a nine-point Self-Assessment Manikin scale^88^. Participants’ heads were stabilized throughout the session with a padded head support frame with an adjustable chin and forehead rest (SR Research Ltd.).

#### Pupil, skin conductance, and facial recordings

Changes in pupil size, skin conductance, and facial expressions were recorded during each of the 60 trials of sound tokens selected from the IADS corpus. Participants sat in isoluminant conditions and fixated on a cross in the center of a front-facing monitor. Sound-evoked changes in pupil size were recorded using the EyeLink 1000 Plus (SR Research Ltd.) at a sampling rate of 1 kHz. Prior to testing, the participant’s pupil size dynamic range was characterized by presenting alternating bright and dark screens on the computer monitor. Skin conductance responses (SCRs) were recorded using the skin conductance module of the BioSemi ActiveTwo system (Cortech Limited Inc.) with two electrodes placed on the hand. Videos of the participant’s facial expressions were recorded at a sampling rate of 120 Hz with a Genie Nano-M2020 camera (Teledyne DALSA) fitted with a 16mm IP/CCTV lens (TAMRON).

#### Mulit-talker speech intelligibility task

The details of the task procedure are similar to previous work^89^. Briefly, participants were required to report four digits that were spoken by a male target speaker (F_0_=115 Hz) masked by two additional speakers (F_0_=90 Hz, F_0_=175 Hz) who were also vocalizing digits. Digits (1-9, excluding the bisyllabic ‘se-ven’) were pseudorandomly selected for each speaker such that each speaker produced a distinct digit at any given time. Stimuli were presented diotically through calibrated circumaural headphones (Bose AE2). After familiarization with the task, participants performed randomly interleaved blocks where four blocks had a signal-to-noise ratio of 9 dB SNR, and four blocks at 0 dB SNR. A block consisted of 10 trials (each a 4-digit sequence), where the first two trials were adaptive in difficulty, designed to re-familiarize the participant with the target, and were excluded from analyses. In each trial, digits were spaced with 0.68 s between onsets, and a virtual keypad appeared 1 s following the fourth digit to allow participants to report the target digits. Feedback was not provided during testing. Pupil size was simultaneously recorded throughout, as described above.

### Quantification and statistical analysis

#### Ranked valence rating

Each participant’s rank-ordered valence ratings were used to order their individual autonomic responses (e.g., in **Figure 2E**). As multiple ratings could have the same integer value, and hence their relative rank-order was arbitrary, all responses for a given integer valence rating were represented by their mean.

#### Token spectrograms

Sounds were filtered via a 64 channel gammatone filterbank with center frequencies spaced between .1 and 10 kHz on an ERB scale. Energy was calculated using 50 ms windows with 25 ms overlap. Values were converted to dB prior to display. Spectrogram plots omit sound frequencies and include additional modifications not present in the source material to honor the terms of use associated with the IADS and IADS-E stimulus batteries.

#### EEG data analysis

Analysis of EEG data was performed in MATLAB (MathWorks, Natick, MA) using FieldTrip software^90^. Data were down-sampled and filtered using zero-phase Butterworth bandpass filters. Eye movement artifacts were identified and removed using independent component analysis^91,92^. The 160 sweeps were averaged together and principal component analysis was used to identify the optimal subset of EEG channels across which to analyze the EFR response^93–95^. After pre-processing, amplitude-intensity functions were quantified using spectrograms, wherein multiple short-term Fourier transforms were computed on consecutive overlapping 1-second intervals using a rectangular window^68^. The second half of the sweep was time-reversed and vector-averaged with the first half of the sweep, combining data from the upward and downward sweeps into a single upward function. A first order polynomial was fit (least-squares) to each growth function between 40 and 65 dB SL. Central gain was quantified as the slope of this fit.

#### Pupillometry data analysis

Analysis of pupillometry data was performed in MATLAB (MathWorks, Natick, MA). To avoid including periods with blinks or missing data, a custom script thresholded absolute pupil size and pupil derivative, padding flagged periods with 100 samples (**Figure S3B**).

Thresholding was verified by visual inspection. For each participant, z-score normalization was performed using the mean and standard deviation pooled from traces in the 3 seconds prior to all 60 trials (**Figure S3C**). The covariate of baseline pupil-size was regressed out linearly (evoked=0.58×baseline+0.39) (**Figure S3D**). Missing data resulting from blink extraction were replaced through linear interpolation. Trials missing more than 50% of data were excluded from analysis Otherwise, flagged missing data were linearly interpolated (**Figure S3E**). Evoked pupil responses were summarized as the mean response between 2 and 5 seconds re. stimulus onset. For the dynamic light stimulus and multi-talker speech intelligibility task, pupillometry analysis procedures matched those described for the responses to IADS stimuli. Trial responses in the multi-talker speech intelligibility task were summarized as the mean pupil diameter between 0.5 and 3.5 second after the onset of the first digit.

#### Skin conductance data analysis

SCR data were pre-processed in MATLAB (MathWorks, Natick, MA) using the Ledalab Version 3.2.2 toolbox^96,97^. A non-negative deconvolution approach was used to separate the skin conductance data into continuous tonic signals (i.e., slow-varying skin conductance level) and phasic signals (i.e., fast-varying SCRs)^96^. For each stimulus trial, the integrated SCR (iSCR) was calculated by taking the time integral of the phasic signal during the eleven seconds following sound onset. The trial average iSCR was calculated for each participant for each sound token.

#### Facial videography data analysis (Figure 3B)

We identified and mapped 478 3-dimensional facial landmarks using the Python MediaPipe toolbox at a downsampled rate of 20 Hz^98^. Face-mesh landmark positions were linearly interpolated across blinks. Landmark positions were temporally smoothed with a Gaussian kernel (sd of 5 samples). Contrast limited adaptive histogram equalization was applied to each video frame. Histogram of oriented gradients analyses (resolution of 8 orientations, local normalization, 2x2 blocks) were performed on each frame, on windows anchored to 77 uniformly spaced landmark. Windows were 20% the height and width of the fitted face-mesh. Euclidean distance relative to features in the preceding/proceeding second was derived and smoothed with a Gaussian kernel (sd of 10 samples). For each participant, Euclidean distances were numerically standardized (i.e., mode and sd calculated with kernel density estimates) across all 60 sound token responses. Trials that deviated by more than five times the average deviation were removed. Facial landmarks were clustered via kernel k-means^99^, where kernels were formed for each participant by calculating the Euclidian distance between facial region responses and applying a Gaussian kernel, before averaging the matrix across participants, and running k-means with 6 clusters (the maximum number of clusters that retained facial symmetry). Trial responses were summarized as the mean across all 77 facial regions between 0 and 6 seconds re. stimulus onset.

#### Elastic net regression

Elastic net regression^100^ was used to investigate the relationship between predictor variables (pupil, face, valence rating, age, and EFR slope), and severity scores (THI, HQ). Predictor variables were first standardized such that fitted coefficients approximated relative predictor strength. Elastic net regression incorporates both regularization and feature selection. The predictive accuracy of the model is penalized for any non-zero coefficients within the model. To regulate correlated predictor variables, the coefficient penalty is a weighted sum of the L1 and L2 norms (here, equally weighted). The value λ scales the size of the coefficient penalty where for larger values of λ any coefficients that are not predictive of the outcome variable are suppressed (**Figure 4A,D**). Fivefold-validation, repeated for 500 random initializations, was used to derive λ that minimized out-of-sample mean square error (MSE). With the predictor variables selected via the elastic net, we refit a linear regression model on all available data (including individuals not in the original elastic net regression due to missing values when including other predictor variables) and contrasted against a linear regression model without the selected autonomic measures (pupil, face) with a likelihood ratio test.

#### Statistical analysis

Statistical analyses were performed with MATLAB (Mathworks, Natick, MA). Non-parametric tests were used when assumptions of parametric tests did not apply (i.e., for behavioral interval data). To assess statistical significance, we used a p-value criterion of p<0.05 (symbolized with an asterisk, when appropriate p<0.01 was also symbolized with two asterisks). Specific statistical details can be found in the corresponding figure legends.

## Notes

### Competing Interest Statement

The authors have declared no competing interest.

### Summary of Updates

Revised abstract. Additional references.

