## Supplemental figures for "The human pupil and face encode sound affect and provide objective signatures of tinnitus and auditory hypersensitivity disorders"

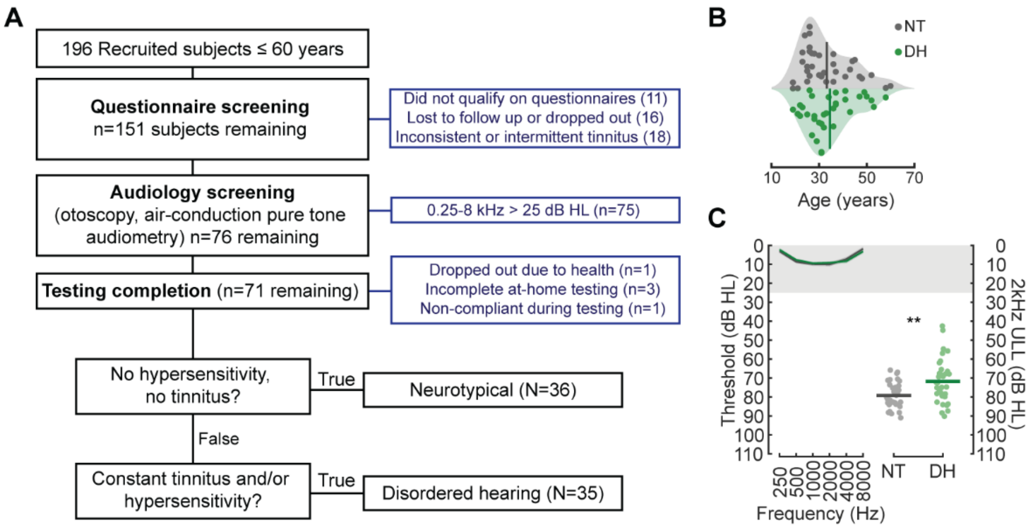


**Supplemental Figure 1** - Participants with disordered hearing are matched for age and hearing thresholds but have lower thresholds for sound discomfort.

**A)** Recruitment and refinement of NT and DH participant cohort.

**B)** Split density functions of participant age. Circles denote individual participants and the vertical line denotes the sample mean (N = 36/35 NT/DH).

**C)** *Left:* NT and DH participants have normal and closely matched hearing thresholds. Thresholds are expressed as dB relative to normal hearing thresholds for each frequency (dB HL). *Right:* The minimum uncomfortable listening level (ULL) at 2 kHz is significantly lower for DH participant (unpaired t-test, $p = 0.0021$). Circles and horizontal bars denote individual participants and sample means, respectively.

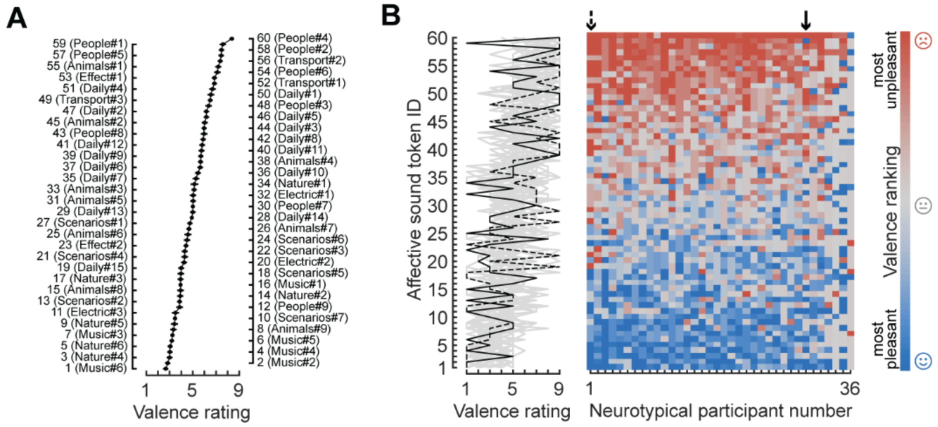


**<u>Supplemental Figure 2</u>** - <u>Participants varied in their selection of sounds that were rated as most pleasant and unpleasant.</u>

**A)** Mean ± SEM rating for the 60 sounds selected from the IADS corpus, in ranked order (N = 36 NT participants).

**B)** Individual valence ratings for 36 NT participants. Solid and dashed lines (left) and corresponding arrows (right) highlight two participants with distinct valence reporting.

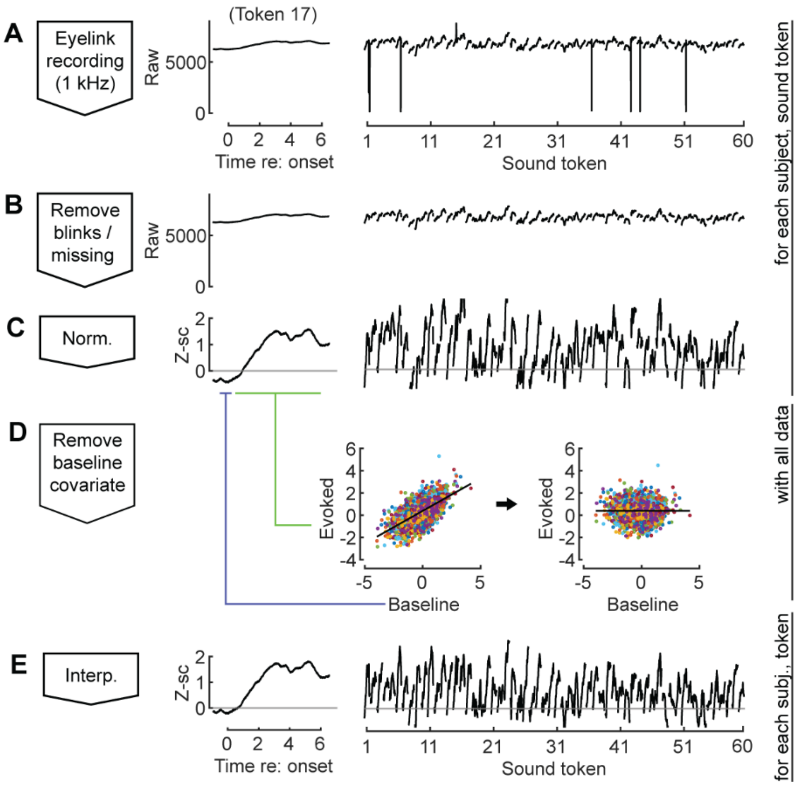


**Supplemental Figure 3 -** Processing of pupillometry data.

**A)** A raw trace recorded via the Eyelink 1000 Plus, from a representative participant listening to the IADS corpus. *Left*, trace in response to a single token is shown. *Right,* responses to all 60 tokens.

**B)** Thresholding was used to identify and remove blinks and missing data.

**C)** To ensure pupil traces were comparable across participants, pupil diameter preceding each IADS token was used to perform z-score normalization.

**D)** Following these preprocessing steps, the linear relationship across all data between baseline pupil size and evoked pupil size was accounted for and removed.

**E)** Missing time samples were filled in through linear interpolation to ensure a continuous trace.

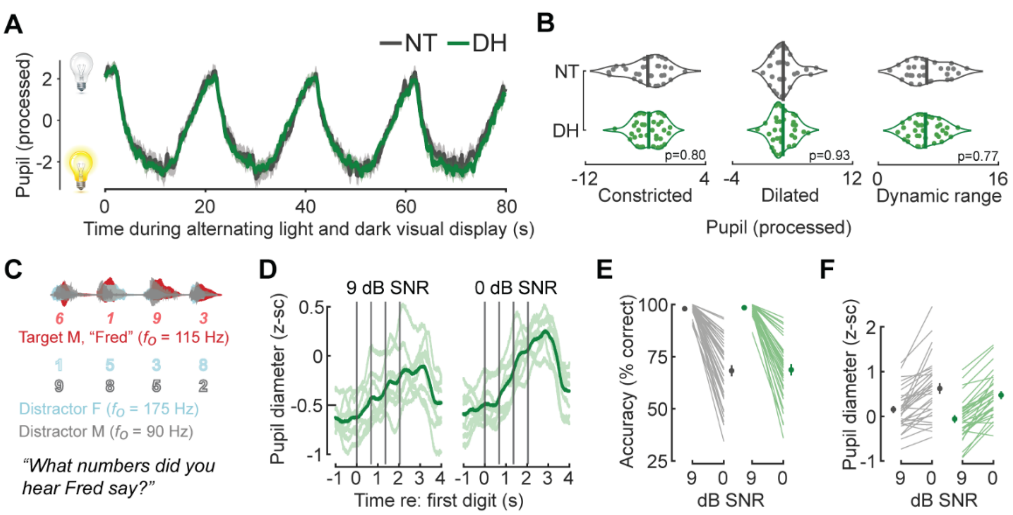


**Supplemental Figure 4 -** Pupil hyper-responsivity is specific to affective processing.

**A)** Pupillary light response measurements were performed to benchmark pupil size in a non-cognitive context. Participants were presented with alternating bright and dark screens from which constricted and dilated states of the pupil could be respectively derived.

**B)** The processed light range pupil response did not differ between NT and DH participants when constricted (left), dilated (middle), or by dynamic range (right).

**C)** Schematic of a multi-talker digit recognition task. Participants (N=36/35 for NT/DH) were familiarized with a target male speaker (red) producing four digits between 1 and 9 (excluding the bi-syllabic '7'), while two spatially co-localized distractors, one male and one female, with $F_0$ frequencies above and below the target speaker simultaneously spoke 4 digits at varying signal-to-noise ratios (SNRs).

**D)** Rapid growth in pupil diameter at the onset of effortful listening is larger overall in more challenging signal to noise ratios. Gray lines denote the timing of each digit in the sequence. Lighter green lines present pupil response for a representative participant across blocks. Thick green line denotes the participant's mean.

**E)** Behavioral accuracy for each individual DH and NT participant and sample mean ± SEM for easy (9 dB SNR) and harder (0 dB SNR) listening conditions. Accuracy is defined as proportion of individual digits correctly reported and is significantly lower in more challenging listening conditions but differences were not observed between groups (Mixed model ANOVA, main effect for SNR [F = 233.4, p = 6 x $10^{-31}$]; main effect for Group [F = 0.06, p = 0.8]; SNR x Group interaction [F = 0.0003 p = 0.96].

**F)** Normalized pupil diameter is significantly larger in more difficult listening conditions, but significant differences were not observed between groups (Mixed model ANOVA, main effect for SNR [F = 28.66, p = 6 x $10^{-7}$]; main effect for Group F = 3.75, p = 0.06]; SNR x Group interaction [F = 0.12, p = 0.73].
